# Fine Motor Serious Game Training Improves Gait in Parkinson’s Disease: A Pilot Study

**DOI:** 10.1101/2025.09.16.676299

**Authors:** Valentin Bégel, Frédéric Puyjarinet, Christian Gény, Valérie Cochen De Cock, Serge Pinto, Simone Dalla Bella

## Abstract

Crucial functions for human behavior such as gross motor skills (e.g., walking), cognitive, rhythmic, and fine motor processes are mostly considered unrelated. However, these functions may interact, but their relations are still poorly understood. Moreover, evidence of causal links between them is scarce. Neurological disorders such as Parkinson’s Disease affect all these functions and thus provide a model to study the interplay between them. We tested the effect of a training delivered using serious games on tablet, involving upper limb rhythmic and fine motor functions, on walking capacities in patients with Parkinson’s Disease (PwPD). PwPD gathered into an Intervention group played either a rhythm game, or an adaptation of *Tetris*, four times a week for six weeks. Before and after the training, gait was evaluated in spontaneous walking and a dual task (counting backward while walking). A Control group of participants did not receive any training. Gait speed, stride length and cadence improved in the Intervention group after the training in comparison with the Control group. The improvement was observed in both Intervention groups, in the spontaneous and in the dual-task conditions. These findings support the hypothesis that gross (axial) motor functions can be trained by fine (lateralized) motor training administered via serious games, possibly by stimulating the rhythmic, perceptual, and cognitive resources sensory systems. This opening promising perspectives for telerehabilitation. Patients with movement disorders could benefit from this promising low-cost and engaging training method, fostering inclusivity and autonomy for people who have reduced access to the clinic.

## Introduction

Theories such as embodied cognition[1,2] and active sensing,[3] as well empirical behavioral and neurofunctional evidence, emphasize the interplay between the cognitive, rhythmic, and fine motor domains and gross motor functions.[4–6] The interrelationship between fine and gross motor domains has been an object of study for decades.[7] Although fine and gross motor domains are theoretically and functionally separated, links between them are consistently reported.[8–10] For example, there is evidence that training gross motor functions (e.g., run, hop, jump) can improve fine motor functions (strike, dribble, kick, catch, throw) in children.[11] However, the opposite causal relationship, namely the effect of fine motor functions training on gross motor functions, remains largely unexplored.

Further evidence of the link between fine and gross motor domains comes from their relationship with cognitive abilities. For example, early locomotor experiences are essential for the development of cognitive skills, and gross and fine motor activities help foster language and cognitive development.[12,13] Cognition and gross motor functions (e.g., gait) are linked in aging, as cognitive skills such as executive function and attention play a pivotal role in motor control.[4,14]

An effective means to address the causal relationship between related cognitive or motor domains is to test patient populations who display deficits in these functions. Parkinson’s Disease (PD) is an interesting model to study the interaction between fine and gross motor domains. People with PD (PwPD) experience fine and gross motor difficulties both in axial (gait, speech, posture) and lateralized (limbs) movements due to the cardinal symptoms associated with the disease (bradykinesia, rigidity, tremor, and postural instability).[15] Fine motor functions are more impaired in PwPD with mild cognitive impairment, compared with patients without cognitive impairment.[16] Training gross motor functions via physical exercise improves cognition as well as gait in PD.[17] This suggests that fine and gross motor domains are related to cognition in PD, and transfer from one domain to another might be observed.

A key aspect of fine and gross motor functions is that sequences of motor events are organized in time. Specifically, activities such as walking or speech articulation, both typically altered in PwPD, rely on rhythmic sequences of movements which are precisely timed.[6,18] The capacities to produce motor rhythms are often assessed with tasks relying on fine motor skills, in tasks such as finger tapping.[19–21] PwPD typically display a more variable motor output in finger tapping tasks.[22,23] Increased variability in PD is also visible in rhythmic tasks involving gross motor functions (e.g., walking,[5]) or in speech.[6,24] Cognition and rhythmic skills are also related, as perceptual and rhythm processing correlate positively with executive functions (e.g., flexibility, inhibitory control), memory, and attention in various population, including PwPD.[25–27]

There is evidence that training gross motor functions can lead to transfer effects in the cognitive, fine and gross motor, and rhythmic domains. Training locomotor functions can improve fine motor skills in children,[11] and cognition in PD.[17] Gait training using rhythmic auditory stimulation improved fine motor and perceptual rhythmic skills (finger tapping with a periodic sound, duration discrimination and detection of misaligned beats in musical excerpts) in PD.[23] However, there is a paucity of studies in which fine motor functions are trained and transfer effects in gross motor functions are tested in PD. To our knowledge, the only exception is a recent study from our group in which fine motor functions were trained with a finger tapping game in PD, and the transfer effects on gait, tapping, and oromotor activity.[28] The focus of this study was narrow, though, as it took motor variability across different motor domains (manual, oromotor, and gait) as the main outcome measure. While the study showed a cross-domain transfer, by demonstrating a reduction in motor variability in the oromotor rhythmic task after the finger tapping rhythmic training, no effect on stride time variability was observed. However, this study did not test other gait spatio-temporal features (speed, stride length, and cadence).

The goal of this pilot study was to assess the effect of serious video game training protocols (one rhythmic game, and one non-rhythmic game) implemented on tablet and requiring finger movements, to improve gross, axial motor functions, namely gait spatio-temporal features, in PwPD. Serious games are games with pleasant and recreational aspects designed for purposes beyond entertainment,[29] and have been proven efficient to train both fine and gross motor domains in PD.[28,30]

We used two games that involve similar finger movements. One of them, *Rhythm Workers*, is a rhythmic game that we developed and validated in PwPD.[21,28,31,32] Notably, this form of rhythmic training differs from standard rhythmic auditory cueing, in which the stimulation is typically delivered while patients walk.[22,23] The other game was an adaptation of *Tetris*, a game that focuses on visuospatial ability and cognitive skills such as decision-making.[33,34] One group of PwPD played *Rhythm Workers* and another played *Tetris* for two months. A third group of PwPD did not receive any training during these two months. Before and after the training period, we tested participants’ gross motor skills (i.e., gait), as well as rhythm perception and cognitive abilities. Some of the participants and the training protocol were the same as in our previously published study;[28] here we focused on the effect of the training on gait spatio-temporal parameters. We predicted that a telerehabilitation training targeting fine motor skills, displayed via serious games (*Rhythm Workers* and/or *Tetris*) would improve participants’ gross motor control involved in gait, compared to a control group who did not receive any training. We also expected that adding a rhythmic component to the training would produce greater transfer to gait performance than non-rhythmic training.

## Methods

### Participants

Participants were recruited by a clinical research assistant, supervised by the team’s neurologist, at the Montpellier University Hospital. We recruited 45 participants with PD. Other outcome measures collected for 33 of these participants were included in a previous publication, in which we focused on speech and motor variability.[28] In this study, we present different gait measures for them and 12 additional participants were tested who received the same training. All participants who were contacted accepted to participate in the study, but three of them withdrew their participation during the experiment, resulting in a final sample of 42 participants (12 women, mean age = 67.55, *SD* = 7.35, age range = 50-82). Patients’ demographic and clinical characteristics are presented in table 1. Patients’ medication was stable for at least 4 weeks prior to the start of the experiment, and they were under optimal medication for the evaluation sessions (60–90 min after their usual dose). Participants were diagnosed with PD in accordance with the UK Brain Bank criteria[15] and disease duration ranged from 2 to 25 years prior to the experiment (mean duration: 10.05 years, *SD* = 5.61 years).

**Table 1.** Patients’ demographic and clinical characteristics.

| Table 1. Patients' demographic and clinical characteristics |  |  |  |
| --- | --- | --- | --- |
|  | Rhythm Workers<br>(n = 19, 3 women) | Tetris<br>(n = 11, 2 women) | No intervention<br>(n = 12, 6 women) |
|  | Mean (range) | Mean (range) | Mean (range) |
| **Demographics** |  |  |  |
| Age | 65.7 (50-77) | 67.7 (56-82) | 70.3 (63-79) |
| Disease duration (Years) | 10.2 (2-25) | 9.6 (2-23) | 10.3 (6-18) |
| **Clinical characteristics (MDS-UPDRS)** |  |  |  |
| Total score | 64.8 (32-102) | 57.4 (24-88) | 56.7 (34-79) |
| Motor subscore (part 3) | 30 (3-57) | 26.3 (10-39) | 29.45 (17-46) |
| Speech item (3.1) | 1.5 (0-2) | 1 (0-2) | 1.2 (0-3) |
| Upper limb items (3.4a–3.6b) | 7.1 (1-16) | 4.7 (0-10) | 8.2 (1-14) |
| Lower limb items (3.7a–3.8b) | 6.5 (0-13) | 5.2 (2-9) | 6.36 (0-12) |
| **Hoehn and Yahr score** | 2.23 (2-3) | 2.2 (2-3) | 2.27 (2-3) |

### Sample Size

Following recommendations by Whitehead et al. (2015),[35] the sample size was estimated without formal power calculation, as it is not required to fulfil the objectives of a small-scale, pilot study. We included a number of participants equivalent to the ones of intervention studies using the same primary outcomes (gait parameters), which corresponds to 10 to 15 participants per group.[22,23]

### Training protocol

Following a parallel-group design, participants were randomly assigned by a clinical research assistant (using the free online application Group Maker - https://livecloud.online/en/group-maker) to the Intervention (n = 30) or the Control group (n = 12). In the intervention group, participants were randomly assigned to the *Rhythm Workers* (*RW* Group, n = 19) or *Tetris* Intervention group (*Tetris* Group, n = 11). Patients in the Intervention group were instructed to play games on a tablet device (LG© G Pad 8.0 model) for at least 30 min, 4 times a week, during a period of 6 weeks, for a total of 12 hours.

*Rhythm Workers* is a music-based rhythm game consisting in finger tapping to the beat of auditory stimuli varying in terms of rhythmic complexity (metronome, metrical sequences, and musical excerpts).[21,32] The goal of the game is to synchronize the finger taps to the beat of rhythmic stimuli. The greater the synchronization (i.e., alignment of the taps with the beats), the higher the score. Feedback is provided by the score (maximum score = 100), and a minimum score (= 70) is required to progress through difficulty levels. *Tetris* is a classical video game in which pieces with different shapes move from the top to the bottom of the computer screen. In the adapted version we developed, the player uses the tactile screen to move and/or rotate the pieces by finger tapping anywhere on the screen, so each piece can fit in with previous pieces to form complete rows (no gaps between pieces). Completed rows disappear, and the remaining pieces drop down. Each completed row earns points, and the level stops when too many pieces filled the screen.

### Performance measures

Performance measures were collected before and after the training at the Montpellier University Hospital by a team of researchers and clinical research assistants from March 2016 to January 2022.

#### Gait

Gait performance was quantified while participants walked over an eight-meter GAITRite© 301 instrumented walkway system (CIR Systems Inc, Havertown, Pennsylvania) composed of pressure-sensitive sensors that activated on foot contact. The walkway was disposed in an isolated hallway free from distraction. Participants started walking two meters away from the walkway to avoid variability (accelerations and decelerations) at the onset of the gait trial. They had to continue walking when they reached the walkway. In a first condition (Spontaneous gait), participants were instructed to walk naturally on the walkway. In a Dual task condition, participants had to verbally count backward from 100, subtracting in increments of seven. The Dual task condition was included to modulate the resource demands attributed to gait, introducing a cognitive load to and stimulate the interplay between gross motor functions and cognition. Three trials were performed in each condition. Spatiotemporal gait parameters, averaged from the three trials, were calculated for both Spontaneous and Dual tasks. Those included Speed (cm/sec), Cadence (steps/minute) and Stride length (cm).

#### Rhythm perception: Beat Alignment Test (BAT)

To control for rhythm perception skills of PwPD, the Beat Alignment Test (BAT) from the BAASTA was administered.[36,37] Participants listened to 72 twenty-beat excerpts from Bach’s Badinerie and Rossini’s William Tell Overture presented at three different tempi (inter-beat intervals, IBIs, of 450, 600 and 750 ms). Starting from the 7th beat, an isochronous sequence with a triangle timbre was superimposed to the music, either aligned (24 trials) to the musical beat or non-aligned (48 trials). Period changes (10% slower or faster than the IBIs, 24 trials) and phase changes (the tones preceded or followed the beats by 33 % of the IBIs, 24 trials) were presented. Participants were asked to judge whether the metronome was aligned or not with the musical beat. The sensitivity index (d’), corresponding to the standardized difference between the hits (i.e., when a misaligned metronome was correctly detected) and false alarms (i.e., when a misalignment was erroneously reported) was calculated as an unbiased measure of detection performance.

#### Neuropsychological assessment

The Montreal Cognitive Assessment[38] was administered to assess global cognitive abilities. Further evaluation of memory and executive functions was performed using the Stroop Color Word Test (interference - time in seconds and number of errors),[39] the Digit Span subtest from Wechsler Adult Intelligence Scale version III (Forward and backward digit span),[40] and the Trail Making Test (TMT, part B, time in seconds and number of errors).[41]

#### Statistical analyses

All analyses were run in R Statistical Software.[42] Analyses of variance (ANOVA) were run with Afex package (v1.0-1)[43] and Tukey-corrected linear contrasts were run with the Emmeans package (1.7-2)[44]. First, we assessed the effect of the training by comparing the Intervention and Control groups. Gait spatiotemporal parameters (speed, stride length and cadence), results at the BAT and neuropsychological tests were entered as dependent variables in a 2-way ANOVA with Group (Intervention vs. Control) as the between-subject factor and Session (Pre vs. Post training) as the within-subject factor.

Z-scores of patients in the Interventions group were calculated relative to the performance of the Control group [Z-score = (*Value - Mean_Controls_)/SD_Controls_*] to ensure that the effect observed in the Intervention group was not merely a test-retest effect. Z-scores were computed across the two types of interventions for gait variables and the BAT scores separately for pre and post-training measures. To quantify the effect of the interventions relative to the performance of the Control group, Delta Z-scores (Δ = *Z-score_Post-Training_* -*Z-score_Pre-Training_*) were calculated for each patient. Positive Delta scores reflect an improvement of the performance. Delta Z-scores for gait spatiotemporal parameters (Speed, Stride length and Cadence) for both the *Rhythm Workers* and the *Tetris* groups were entered as dependent variables in a 2-way ANOVA with Group (*Rhythm Workers* vs. *Tetris*) as the between-subject factor and Condition (Spontaneous vs. Dual task) as the within-subjects factor. General Eta Squared are reported as a measure of effect sizes (.01: Small effect size, .06: Medium effect size,14 or higher: Large effect size). In addition, the spontaneous and Dual-task conditions were averaged, and t-tests were used to test if the mean of the Delta Z-scores at the gait parameters differed from zero (absence of an effect of the training) in each group.

Finally, we used the clinically meaningful criteria in PD defined by Hass and collaborators to quantify the benefits of the training at the individual level.[22,45,46] Following Hass and collaborators criteria, a difference of 0.06 m/s in gait speed was considered as small, a difference of 0.14 m/s was moderate, and a difference of 0.22 m/s was considered as large. We then divided participants into two categories based on their response to the training. Participants who showed a clinically significant improvement in gait speed (at least 0.6m/s in either Spontaneous or Dual task conditions) were considered as participants with a Positive Response (PR). Participants who did not respond positively to the training were considered as participants with a Non-Positive Response (NPR, for a similar classification of patients’ response to a rhythmic training see Dalla Bella et al., 2017[22]). The difference between the PR and NPR groups in total time spent playing the game (in hours) was assessed using Wilcoxon rank-sum test. We computed t-tests on the Delta Z-scores to compare the effect of the training on gait Speed, Stride length and Cadence in the PR and NPR groups. For each variable, significance values were corrected for multiple comparisons.

## Results

### Intervention vs. Control

Gait spatio-temporal parameters (speed, stride length, and cadence) at Pre-training and Post-training evaluation sessions are shown in Figure 1. Notably, the Intervention and Control groups did not differ at baseline (Speed, Spontaneous: *p* = .39, Dual task: *p* = .69; Stride length, Spontaneous: *p* = .52, Dual task: *p* = 62; Cadence, Spontaneous: *p* = .49, Dual task: *p* = 46). Overall, participants in the Intervention group increased their gait speed by 5.01cm/second (4.67%) in the Spontaneous task and 10.14cm/second (13.34%) in the Dual task, stride length by 3.93cm (3.24%) in the Spontaneous task and 7.26cm (6.61%) in the Dual task, and cadence by 1.25 step/min (1.18%) in the Spontaneous task and 5.95 step/min (7.33%) in the Dual task from pre to post training. Participants in the Control group showed no significant change.

**Figure 1.**
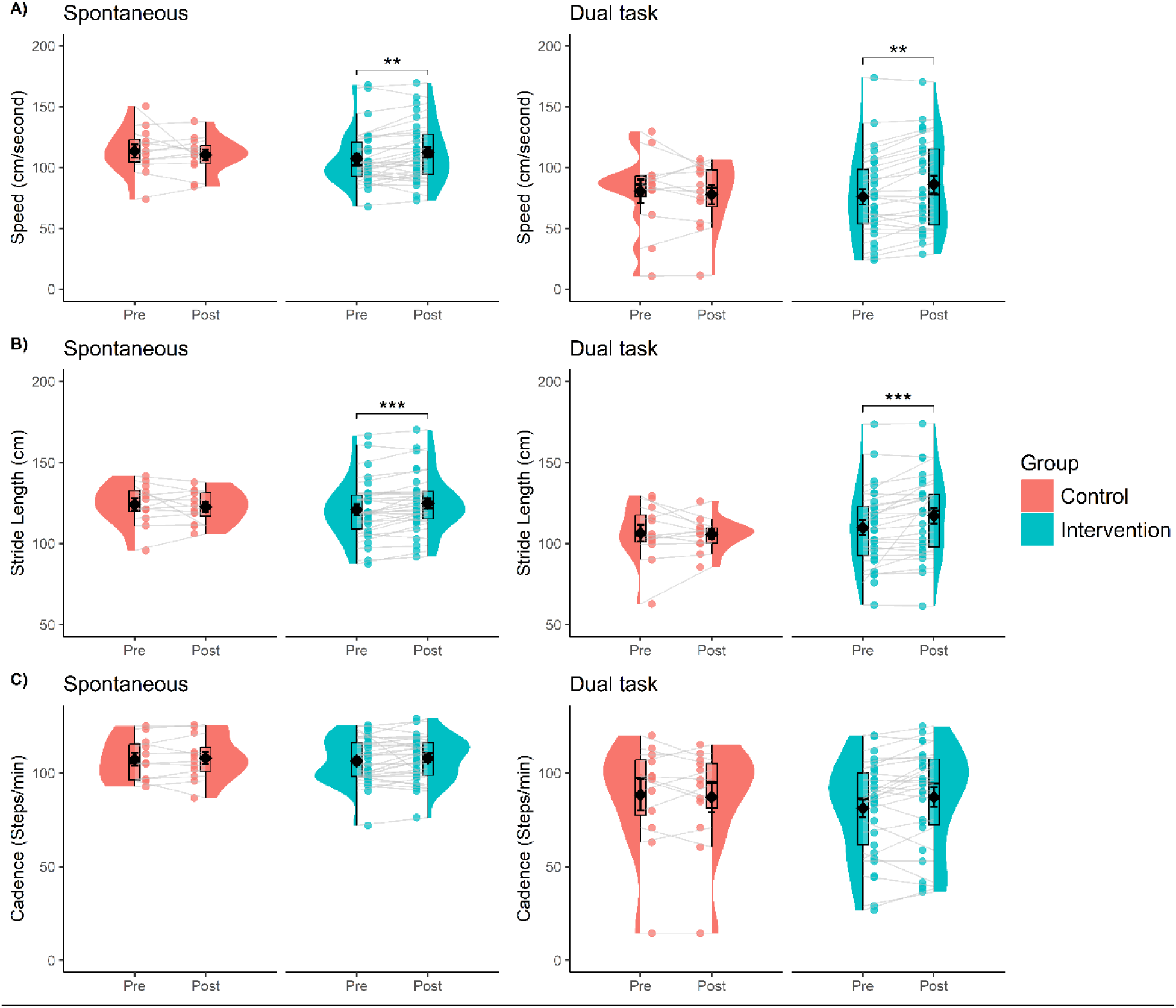
Gait spatiotemporal parameters (A = Speed, B = Stride length, C = Cadence) in the Spontaneous task and Dual task for the Interventions and the Control groups. Errors bars represent Standard Errors of the mean.

For gait Speed, the two-way ANOVA pointed to a Group x Session interaction, at the threshold of statistical significance, *F*(1,40) = 3.90, *p* = 0.055, 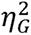 = 0.07 (Figure 1A, left panel). The difference between Pre and Post was significant in the Intervention group, *t*(40) = 2.25, *p* = 0.03, *d* = 0.58, but not in the Control group, *t*(40) = 0.91, *p* = .37, *d* = 0.37. The Group X Session interaction attained significance in the Dual task condition, *F*(1,37) = 6.25, *p* = 0.017, 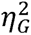 = 0.006 (Figure 1A, right panel). The difference between Pre and Post was significant in the Intervention group, *t*(37) = 3.37, *p* = 0.002, *d* = 0.92, but not in the Control group, *t*(40) = 0.75, *p* = 0.45, *d* = 0.31.

For Stride length, there was a Group x Session significant interaction in the Spontaneous condition, *F*(1,40) = 5.22, *p* = 0.028, 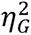 = 0.006 (Figure 1B, left panel). The difference between Pre and Post was significant in the Intervention group, *t*(40) = 3.04, *p* = 0.004, *d* = 0.79, but not in the Control group, *t*(40) = 0.77, *p* = 0.44, *d* = 0.32. In the Dual task condition, there was an interaction between Group and Session on Stride length, at the threshold of statistical significance*, F*(1,37) = 4.01, *p* = 0.053, partial 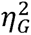 = 0.008 (Figure 1B, right panel). The difference between Pre and Post was significant in the Intervention group, *t*(37) = 3.28, *p* = 0.002, *d* = 0.89, but not in the Control group, *t*(37) = 0.22, *p* = 0.83, *d* = 0.90.

No main effects or interaction were found for Cadence (Spontaneous and Dual task conditions), rhythm perception (The BAT) and in the neuropsychological tests (all *p*s > .05, see supplementary table).

### Rhythm Workers vs. Tetris

The Deltas, measured as the difference between the post and the pre-training at the averaged Spontaneous and Dual tasks Z-scores (relative to the Control group for the *Rhythm Workers* and *Tetris* subgroups) are shown in Figure 2. As can be seen, the Z-scores are generally larger than zero, indicating an improvement of the performance in both groups.

**Figure 2.**
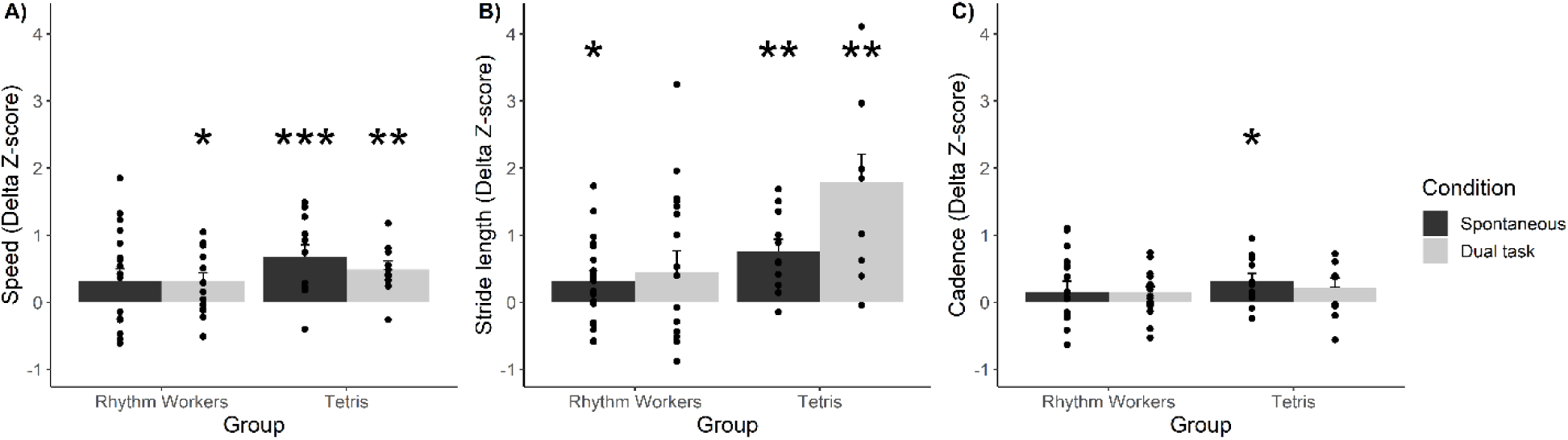
Gait spatiotemporal results (Z-scores) at the gait parameters for the Rhythm Workers and the Tetris groups. Zero on the Y axis means that there is no improvement. * p < .05, ** p < .01, *** p < .005, one-sample t-test against zero (zero = no effect of the interventions), corrected for the four comparisons in each panel using Bonferroni correction. Errors bars represent Standard Errors of the mean.

The ANOVA on gait Speed revealed no significant main effect or interactions.

The ANOVA on Stride length revealed a significant main effect of Group*, F*(1,25) = 6.05, *p* = 0.021, 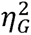 = 0.160, suggesting that in general, the effect was more important in the *Tetris* group. The effect was also more important in the Dual task condition (Main effect of Condition, *F*(1,25) = 8.00, *p* < 0.01, 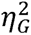 = 0.009). Finally, there was an interaction between Group and Condition factors, *F*(1,25) = 5.63, *p* < 0.026, 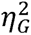 = 0.46. In the *Tetris* group, Delta Z-scores were significantly larger in the Dual task (*M* = 1.79, *SD* = 1.31) than in the Spontaneous (*M* = 0.76, *SD* = 0.59), *t*(25) = 3.27, *p* = 0.003, *d* = 1.47, whereas they did not differ in the RW Group (Spontaneous, *M* = 0.32, *SD* = 0.64; Dual task, *M* = 0.45, *SD* = 1.28), *t*(25) = 0.37, *p* = 0.71, *d* = 0.13.

For Cadence, there was no significant main effect or interactions.

We assessed whether differences in gait improvement were linked with the amount of time participants played the game in the two groups. Participants in the *Tetris* group played significantly more time than in the RW group (RW, *M* = 5h, *SE* = 0.9h; *Tetris*, *M* = 29h, *SE* = 0.9h), *t*(10.23) = 2.81, *p* < 0.018, *d* = -1.40).

Although there were no effects of the interventions on the results at the BAT and the neuropsychological tests, we conducted the Z-score analyses for these variables to highlight the different effects of each training on rhythmic and cognitive skills. Participants in the RW group had larger Delta Z-scores than participants in the *Tetris* group at the BAT (RW, *M* = 0.30, *SE* = 0.19; *Tetris*, *M* = -0.35, *SE* = 0.14), *F*(1,25) = 4.86, *p* = 0.037, 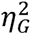 = 0.163. We ran correlations between the Delta Z-scores in the BAT and the gait variables to explore the relationship between the improvement in rhythmic skills and in gait separately for each group. None of the correlations were significant (all *p*s > .05). The two groups did not differ in the neuropsychological tests (all *p*s > .05).

### Interindividual differences

Overall, 20 patients out of 31 (67%) showed a clinically meaningful improvement (PR) in the Intervention group. Eight of them showed a small effect (5 in the RW group, 3 in the Tetris group), six a moderate effect (2 in the RW group, 4 in the Tetris group) and three a large effect (2 in the RW group, 1 in the Tetris group) in Gait speed. The gait performance of PR and NPR in the Intervention group (RW and Tetris groups taken together) are presented in Figure 3. The improvement of gait parameters was significantly more important in PR than in NPR for Speed (PR, *M* = .60, *SE* = .15; NPR, *M* = -.02, *SE* = .11), *t*(16.50) = 3.47, p = .001, *d* = 1.39, Stride length (PR, *M* = .94, *SE* = .19; NPR, *M* = .12, *SE* = .30), *t*(16.56) = 2.31, p = .017, *d* = 0.92, and Cadence (PR, *M* = .43, *SE* = .09; NPR, *M* = -.24, *SE* = .22), *t*(12.24) = 2.85, p = .007, *d* = 1.31.

**Figure 3.**
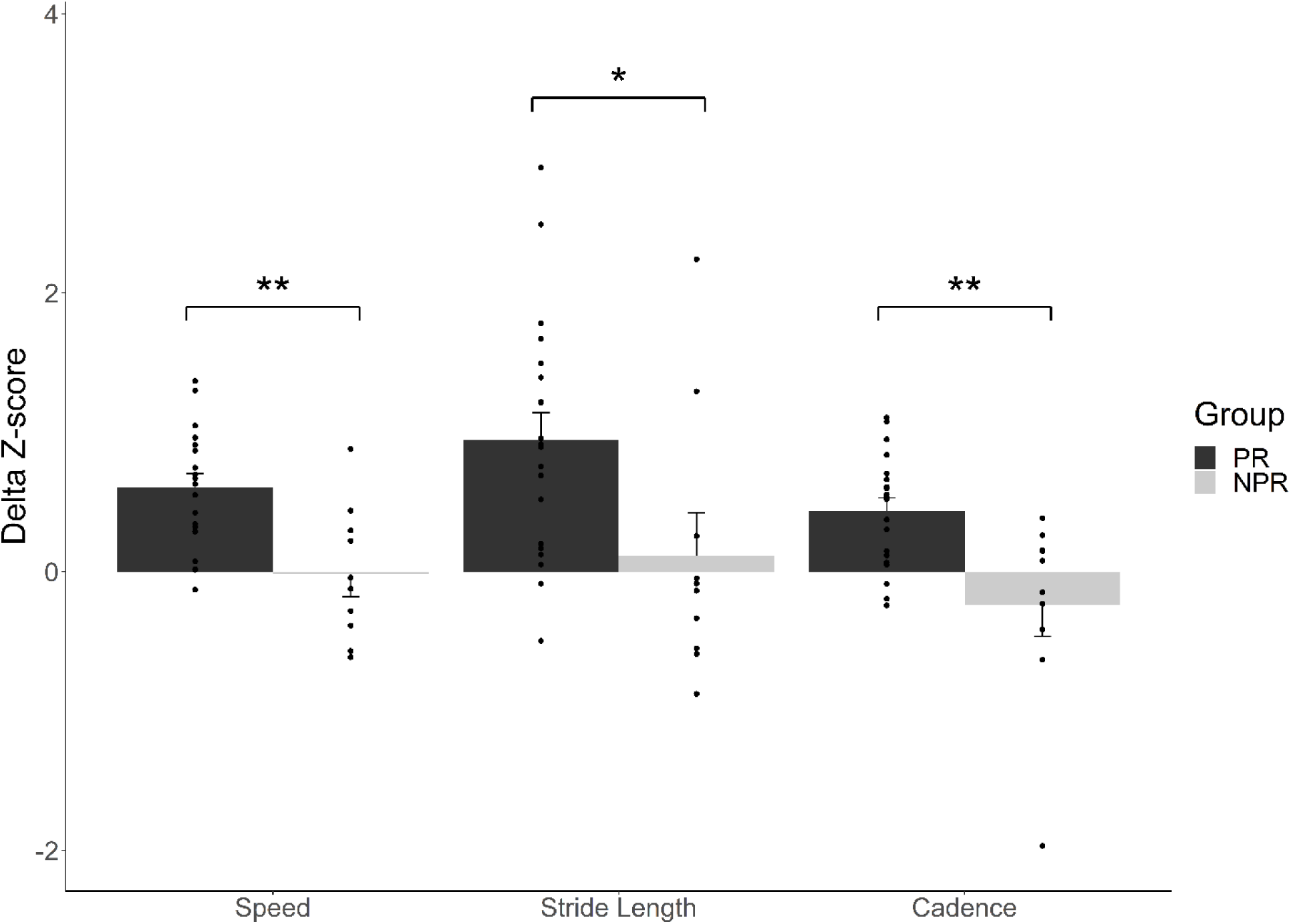
Gait spatiotemporal results (Z-scores for the gait parameters) for PR and NPR. Average values for the Spontaneous and Dual task conditions are presented. * p < .05, ** p < .01

Across the two groups, PR played significantly more than NPR (PR, *M* = 17h, *SE* = 5.5h; NPR, *M* = 7h, *SE* = 1.5h; *t*(21.52) = -1.77, *p* = .045, *d* = 0.49).

## Discussion

The aim of this pilot study was to test the effect of playing mobile video games involving fine motor movement (finger movement) on one gross motor skill (gait). We provided first evidence that spatio-temporal parameters in gait can be trained in patients in PD patients playing a video game on tablet (a rhythm game or an adaptation of *Tetris)* training fine movement abilities for six weeks. After the training period, patients showed improved walking, as revealed mostly by faster speed, and increased stride length. The effect was visible in spontaneous walking (4.67% for speed and 3.24% for stride length) as well as in a dual-task condition consisting in counting backwards while stepping (12.34% for speed and 6.61% for stride length). In comparison, a control group of PD participants without training did not show any improvement.

The observed transfer effect from a fine motor control training, like finger tapping, to gait spatio-temporal parameters is novel. Participants who trained fine motor control - in a rhythmic or non-rhythmic game - displayed improvements of spontaneous gait performance just slightly lower than observed after standard rhythmic auditory cueing therapies, and superior with regard to dual-task performance. For example, in a previous study, PD patients improved their speed by 6% and their stride length by 4.5% in spontaneous walking after they underwent a one-month rhythmic auditory cueing program.[22]. This is consistent with the average improvement observed in recent meta-analyses of the effect of auditory cueing in PwPD.[47,48] Moreover, if we focus on the gait effects of deep brain stimulation protocols, the intervention typically does not show a superior general improvement. Patients who respond positively to deep brain stimulation in terms of gait speed (53%) show an improvement of 5.72% on average.[49] In sum, comparisons with previous studies suggest that fine motor training improves gait parameters in spontaneous walking to a level slightly inferior to the benefits of rhythmic auditory cueing. This is not surprising, as rhythmic auditory cueing directly trains gait, thus within-effector benefits are expected.[22,50–52] Nevertheless, the effect was still quite high in spontaneous walking and even more in dual task, compared for example with deep brain stimulation, given that gait was not trained. Notably, more than two thirds of patients (67%) responded positively to the training by showing a clinically meaningful improvement measured by gait speed, with almost one third (29%) exhibiting average to large improvements.[45] This finding is promising, considering that usually around 50% of the patients typically respond positively to rhythmic auditory cueing (see recent studies that used the same criteria[45] to quantify the clinical benefits of the training protocol[22,46]), which is similar to what is observed after deep brain stimulation.[49]

Moreover, there are important individual differences in the response to the training, which may be partly explained an effect of dose, namely the true amount of training received. Participants who responded positively to the training played on average more (17h) than the instructed time (12h). In contrast, the ones who did not respond to the training played significantly less (7h). In a future clinical trial, efforts should be made to increase participants’ adherence to the training, by improving the game design and add features that are more engaging for PwPD.

The beneficial effects of gamified training of fine motor movements were visible for the rhythmic game and Tetris alike. However, the mechanisms underlying these improvements are likely to be different, as the two games engage different perceptual and cognitive functions. For example, it has been shown in previous studies that that rhythmic auditory cueing is linked to rhythmic sensorimotor skills, making rhythmic processes that are independent from an effector a plausible mechanism underlying the improvement of gait parameters.[22,23,46] In line with this idea, we reported in a recent study that rhythmic training in PD involving finger tapping reduces motor variability in gait.[28] Different mechanisms may underlie the positive effect of *Tetris.* The absence of improvement at a perceptual rhythmic task (the BAT) after the training in the *Tetris* group confirms that *Tetris* does not influence rhythmic skills. The effect of *Tetris* on gross motor skills could be attributed to selective perceptual and cognitive improvements. Balance and gait are complex, biomechanically constrained skills that require perceptual (sensory strategies, orientation in space) and cognitive resources.[53] Previous studies have shown that although *Tetris* does not improve general cognitive skills,[54] it enhances specific skills such as selective visual attention[55] and spatial skills.[56] These skills are impaired in PD and contribute to gait and posture disorders.[57,58] Recently, Wielenski and colleagues conducted a pilot study that showed that training visual attention skills improves gait in PD studies.[59] Sufficient visual attentional and spatial orientation resources are necessary for postural control, and more so when the task is complex, such as in dual-tasks.[60] The beneficial effect of the *Tetris* training on gait is stronger in the dual-task condition (Stride length), further suggesting that *Tetris* training is tapping into visual attention and spatial orientation skills. Although it is possible that the positive effect of *Tetris* and *Rhythm Workers* are based on different mechanisms, we cannot rule out the involvement of a common mechanism, possibly based on reward and motivation induced by playing. Music and video games activate dopaminergic pathways associated with reward processing and motivation.[61,62] Dysfunctions in these dopaminergic pathways are one of the main physiopathological features of PD.[63] Playing video games may therefore improve the activity of the dopaminergic system impaired in PD.

In conclusion, this short-scale study supports the hypothesis that one gross motor skill (gait) can be improved by fine motor, finger-tapping, training administered via video games. Findings in PwPD revealed a possible causal relationship between fine and gross motor skills. A future clinical trial with more participants will be needed to confirm these results, with an improved game design to increase participants’ adherence to the training. Future studies with more exhaustive testing of motor, perceptual and cognitive skills before and after the training’ to further characterize the mechanisms that mediate the transfer effect from fine motor to gross motor skills. This will also serve to rule out a placebo effect, as the effect of the training in the intervention group was shown in comparison with a passive control group. Nevertheless, the fact that positive responders played more than non-positive responders suggests that the training *per se* was efficient.

The results of this study have implications for clinical research and healthcare. Most methods to train gross motor skills require costly equipment as well as the presence of a clinician, and thereby cannot be carried out at home. The telerehabilitation technique that we used is low-cost, user-friendly, and fosters inclusivity and autonomy, as it could provide people who have reduced access to clinics or patients who are temporarily immobile with an efficient training method.

## Statements and Declarations

### Ethics approval and consent to participate

This study was approved by the French Comité de Protection des Personnes (CPP N_2015-A01090-49). All participants gave their informed consent in accordance with the Helsinki Declaration (World Medical Association, 2024). This study was registered as a clinical trial (ClinicalTrials.gov, registration NCT02855710).

### Availability of data and materials

The datasets used and/or analyzed during the current study are available from the corresponding author on reasonable request.

### Competing Interests and Funding

Simone Dalla Bella and Valérie Cochen De Cock are board members of the BeatHealth startup dedicated to the design and commercialization of rhythm-based testing devices (including BAASTA) and cognitive therapies such as Rhythm Workers. For the remaining authors, no competing interests are to be declared.

This research was funded by a CIFRE grant (R153304C) to VB, the Junior Grant from the Institut Universitaire de France (IUF) to SDB, and the Maturation Technological Transfer grant (YouRhythm project) from SATTAxLR to SDB and the company NaturalPad.

### Author contributions

V.B., S.P., and S.D.B conceived and planned the study. V.B, F.P., and C.G. collected the data.

V.B. and S.D.B analyzed, interpreted the data, and drafted the manuscript. All authors revised the paper.

## Acknowledgments

We thank all PwPDs for their collaboration in this study.

---

**Supplementary table.** Scores at the neuropsychological tests and the BAT.

|  |  |  |  |  |
| --- | --- | --- | --- | --- |
|  | Intervention (N = 30) |  | No intervention (N = 12) |  |
|  | Mean (SD) |  | Mean (range) |  |
| Total score | Pre | Post | Pre | Post |
| MoCA | 25.30 (2.73) | 25.42 (2.73) | 25.73 (3.00) | 25.73 (3.00) |
| Stroop (Errors) | 2.76 (3.26) | 3.22 (3.19) | 2.60 (2.78) | 2.95 (1.54) |
| Stroop (Time) | 81.26 (42.76) | 85.80 (53.62) | 87.69 (29.57) | 92.60 (27.13) |
| Digit Span (Forward) | 8.11 (2.15) | 8.00 (1.98) | 8.91 (2.47) | 7.91 (0.35) |
| Digit Span (Backward) | 5.59 (1.80) | 5.32 (1.83) | 5.27 (2.05) | 5.64 (2.42) |
| TMT B (Errors) | 0.78 (0.90) | 0.63 (1.05) | 1.18 (1.15) | 1.05 (1.44) |
| TMT B (Time) | 67.00 (30.22) | 72.46 (46.51) | 85.14 (26.70) | 81.86 (25.55) |
| BAT | 1.79 (1.13) | 1.96 (1.03) | 1.84 (0.85) | 1.87 (0.87) |

## Notes

### Competing Interest Statement

Simone Dalla Bella and Valerie Cochen De Cock are board members of the BeatHealth startup dedicated to the design and commercialization of rhythm-based testing devices (including BAASTA) and cognitive therapies such as Rhythm Workers. For the remaining authors, no competing interests are to be declared.

### Summary of Updates

Abstract revised. Minor changes throughout the manuscript

